# Heimdallarchaeota harness light energy through photosynthesis

**DOI:** 10.1101/2020.02.20.957134

**Authors:** Rui Liu, Ruining Cai, Jing Zhang, Chaomin Sun

## Abstract

Photosynthesis is an ancient process that originated after the origin of life, and has only been found in the Bacterial and Eukaryotic kingdoms, but has never been reported in any member of the domain Archaea. Heimdallarchaeota, a member of Asgard archaea, are supposed as the most probable candidates (to date) for the archaeal protoeukaryote ancestor and might exist in light-exposed habitats during their evolutionary history. Here we describe the discovery that Heimdallarchaeota genomes are enriched for proteins formerly considered specific to photosynthetic apparatus and are suggestive performing oxygenic photosynthesis. Our results provide strong support for hypotheses in which Heimdallarchaeota harvest light by bacteriochlorophyll and/or carotenoid, then transport electron from photosystems to Calvin-Benson-Bassham cycle along with CO_2_ fixation and ATP biosynthesis, and release oxygen as a waste product. Given the possessing of phototrophic lifestyle together with other anaerobic and aerobic metabolic pathways, Heimdallarchaeota are firmly believed to be photomixotrophic and have a facultative aerobic metabolism. Our results expand our knowledge that archaea have played an important role in the molecular evolution of eukaryotic photosynthesis and raise the significant possibility that Heimdallarchaeota might be ancestor of eukaryotic photosynthetic organisms.

## Introduction

Photosynthesis, the foundation for life, result in an enormous increase in biomass production on Earth and produce oxidized compounds serving as electron acceptors for heterotrophic metabolism^1^. In microbiology, organisms that perform photosynthesis include Cyanobacteria, Chloroflexi, Firmicutes, Chlorobi, Proteobacteria, Acidobacteria and Gemmatimonadetes^2^. Among them, only Cyanobacteria carry out oxygenic photosynthesis using electrons originating from water to generate oxygen as product, and evolve in ancestral near the time to the rise of oxygen and eventually result the Great Oxidation Event on Earth^3^. Till date, microbial photosynthesis has been only found in bacteria but not in any reported archaea except a functional bacteriochlorophyll synthase identified in an uncultivated Crenarchaeota, which provides the only clue of photosynthesis existing in Archaea^4^.

Breakthroughs in environmental and metagenomic sequencing technologies are rapidly transforming the landscape for microbial evolution, especially the discovery of Asgard phylum archaea and their supposed position at the base of the eukaryotic tree of life^5^. It stands to reason that many phototrophs remain to be discovered, thus we want to ask if these metagenomic efforts help us to uncover phototrophic Archaea. Heimdallarchaeota are a member of Asgard superphylum archaea and currently represent the predicted closest archaeal relative of eukaryotes, and might exist in light-exposed habitats in their evolution history^5–8^. However, due to far less assembled genomes of Heimdallarchaeota than other Asgard members, no more light-dependent lifestyle details of this uncultured archaea have been disclosed. In the present study, nine high-quality assembled genomes of Heimdallarchaeota were obtained, and specific analyses of their reconstructed genomes provide solid proof that Heimdallarchaeota could utilize light energy through bacteriochlorophyll and/or carotenoid-based oxygenic photosynthesis. Given the closest eocytic lineage to eukaryotes, Heimdallarchaeota are proposed to be ancestor of eukaryotic photosynthetic organisms.

## Results and Discussion

### Photosynthetic apparatus in Heimdallarchaeota

To gain insight into the light utilizing characteristics in Heimdallarchaeota, we sampled aquatic sediments from a typical cold seep in South China Sea and two hydrothermal vents in Western pacific (Expanded Data Fig. 1 and Extended Data Table 1) with distinct chemical parameters (Expanded Data Fig. 2). Total DNA was extracted from all samples and sequenced, and nine high quality (>50% completeness, <10% contamination) metagenome-assembled genomes (MAGs) of Heimdallarchaeota were obtained by utilizing a hybrid binning strategy and performing manual inspection and data curation (Extended Data Table 2). The maximum-likelihood phylogenetic tree, based on concatenation of 37 marker genes including 13 small subunit (SSU) and 16 large subunit (LSU) ribosomal RNA genes, showed that both our and published Heimdallarchaeota MAGs clustered with other Asgard superphylum members and displayed a much closer evolutionary linkage with eukaryotes than other archaeal superphyla including DPANN, TACK and Euryarchaeota (Extended Data Fig. 3 and Supplementary Table 1). In accordance with other Asgard members, credible eukaryote-specific proteins (ESPs) were identified in our Heimdallarchaeota MAGs (Extended Data Fig. 4, Supplementary Table 2), which confirms Heimdallarchaeota as the current best candidate for the closest archaeal relatives of the eukaryotic nuclear lineage as described previously^5,6^. Different with previous report about the existence of rhodopsins in Heimdallarchaeota^1^, there is no any rhodopsin homologs identified in the present nine Heimdallarchaeota MAGs. However, many typical chloroplastic proteins (including protochlorophyllide reductase, chlorophyll(ide) b reductases NOL/NYC1, NAD(P)H quinone oxidoreductase, photosystem I assembly proteins Ycf3 and phycocyanobilin lyase) were surprisingly identified in our Heimdallarchaeota MAGs (Extended Data Fig. 4 and Supplementary Table 2), indicating that Heimdallarchaeota might be a kind of unprecedented photosynthetic organism.

To carefully determine the photosynthetic position of Heimdallarchaeota, we performed various in-depth photosynthetic analyses based on our and published Heimdallarchaeota MAGs. It is well known that for the energy of sunlight to be converted and stored into biological systems, it must first be captured by the pigments present in the photosynthetic organisms. All photosynthetic organisms synthesize two types of pigments: (a) bacteriochlorophylls and/or chlorophylls, which function in both light harvesting and photochemistry, and (b) carotenoids, which primarily act as photoprotective pigments but can also function in light harvesting^9^. Notably, almost all necessary bacteriochlorophyll synthesis components widely distribute in Heimdallarchaeota MAGs, which provides sufficient evidence that Heimdallarchaeota could synthesize bacteriochlorophyll (Fig. 1a, Supplementary Table 3). Among the key enzymes synthesizing bacteriochlorophyll, protochlorophyllide reductase (Por) could catalyze the reaction of transition between divinyl protochlorophyllide and divinyl chlorophyllide a^10^. By the phylogenetic analysis, most of Por homologs in our Heimdallarchaeota MAGs clustered in a single clade at the root of the tree (Fig. 1b, Supplementary Table 3). However, one Por homolog in H2.bin.2 was found to locate in a sister clade with typical photosynthetic organisms including Cyanobacteria, Algae and Streptophytina (Plants), and they further clustered with phototrophic bacteria containing Chlorobi, Chloroflexi and Proteobacteria (Rhodospirillales and Chromatiales), revealing the potential Heimdallarchaeota-phototrophs affiliation. The bacteriochlorophyll synthase (BCS) is capable of synthesizing bacteriochlorophyll a by esterification of bacteriochlorophyllide with phytyl diphosphate or geranylgeranyl diphosphate^4^, and it is also annotated as digeranylgeranylglyceryl phosphate synthase (DGPS). We further phylogenetically analyzed the evolutionary relationship between BCS and DGPS, and the results showed that all archaeal DGPS located at the root, which separating from the clade containing BCS from phototrophic bacteria and chlorophyll synthase from photosynthetic organisms (Fig. 1c, Supplementary Table 3). And homologous proteins of BCS in our Heimdallarchaeota MAGs clustered a clade with the DGPS in Heimdallarchaeota LC2, which located at a position between DGPS and BCS branches (Fig. 1c, Supplementary Table 3). Furthermore, a previous reported functional bacteriochlorophyll synthase derived from the uncultured Crenarchaeota^4^ was found to cluster a branch with the DGPS from Heimdallarchaeota LC3^5^, and this cluster displayed a close evolutionary relationship with those photosynthetic bacteriochlorophyll and chlorophyll synthase branches (Fig. 1c). Although the function of these putative bacteriochlorophyll associated proteins remains to be elucidated, it is tempting to speculate that Heimdallarchaeota have a bacteriochlorophyll producing capability. It is known that bacteriochlorophylls or chlorophylls exist in only photosynthetic organisms^3,11^. The evidence of bacteriochlorophyll production in both Heimdallarchaeota and Crenarchaeota suggests that archaea should have played an important role in the molecular evolution of bacteriochlorophyll synthase, and raises the significant possibility that the origin of photosynthesis probably predates the divergence of bacteria and archaea.

**Fig. 1.**
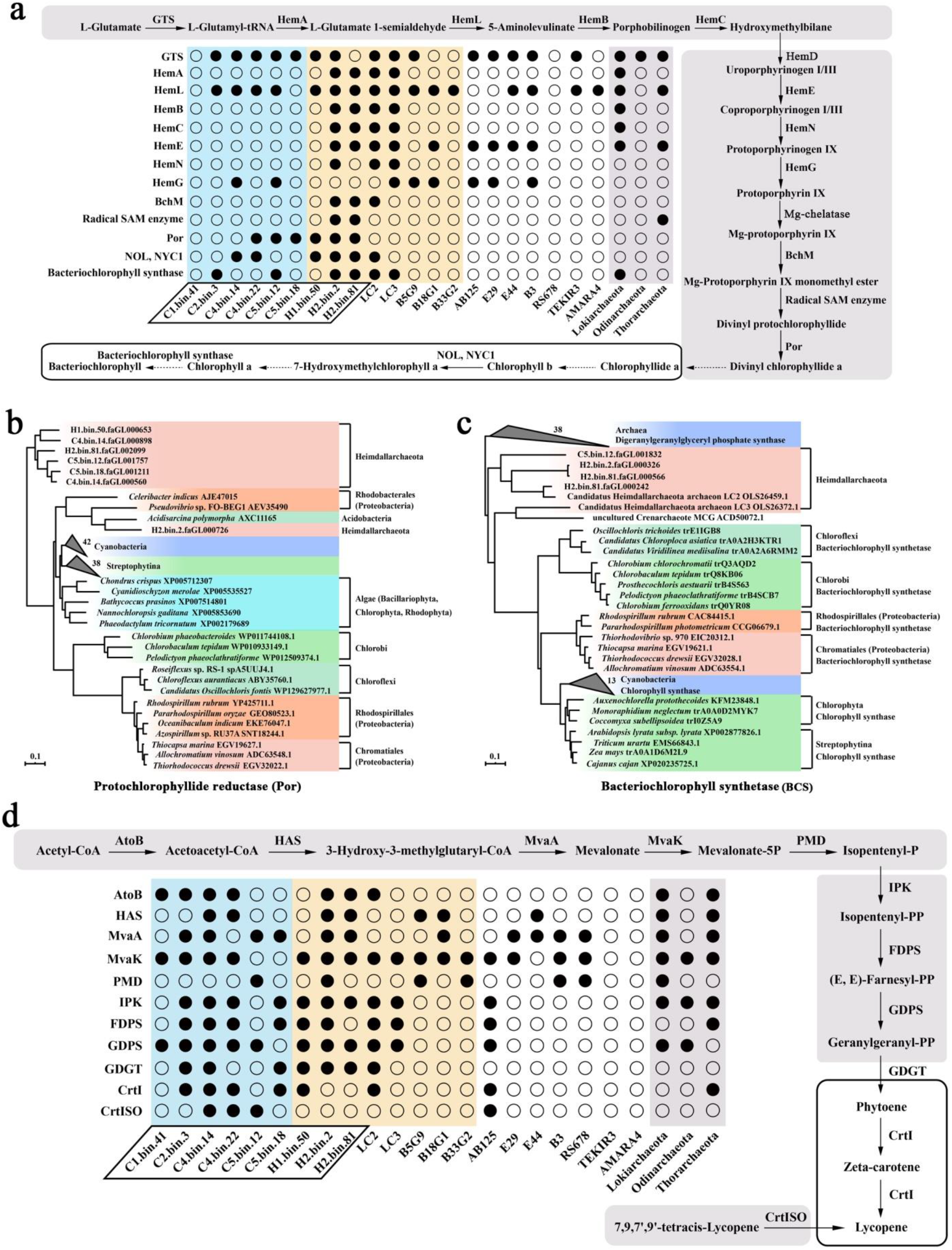
Phylogenomic analysis of photosynthetic pigments biosynthesis in Heimdallarchaeota. **a**, Analysis of bacteriochlorophyll biosynthesis in different Heimdallarchaeota and other Asgard archaeal genomes. **b**, **c**, Phylogenetic analyses of protochlorophyllide reductase (Por) and bacteriochlorophyll synthetase (BCS). A rooted maximum-likelihood tree of Por (b) or BCS (c) homologs derived from different photosynthetic organisms identified in this work. The bootstrap support values 1,000. All proteins and species detailed information used for phylogenetic analyses are listed in Supplementary Table 3. **d**, Analysis of lycopene biosynthesis in different Heimdallarchaeota and other Asgard archaeal genomes. For panels a and d, the solid arrows indicate the enzymes associated with bacteriochlorophyll or lycopene biosynthesis identified in Heimdallarchaeota MAGs. Dotted arrows indicate the enzymes associated with bacteriochlorophyll or lycopene biosynthesis not identified in Heimdallarchaeota MAGs. The light blue box highlights Heimdallarchaeota MAGs from cold seeps. The light pink box highlights Heimdallarchaeota MAGs from hydrothermal environment. The light purple box highlights assembled genomes of other Asgard archaea. The parallelogram box highlights Heimdallarchaeota MAGs obtained in this study. The detail information of key enzymes involved in bacteriochlorophyll and lycopene biosynthesis is listed in Supplementary Tables 3 and 5.

In addition bacteriochlorophyll, other photosynthetic pigments including carotenoid^9^ and bacteriophytochrome^12^ were also identified to be synthesized in Heimdallarchaeota. Particularly for carotenoid, it is a kind of ubiquitous and essential pigment in photosynthesis^9^, and functions as accessory light-harvesting pigment and transfers the absorbed energy to bacteriochlorophylls, which expands the wavelength range of light that is able to drive photosynthesis^13,14^. We reconstructed the complete synthesis pathway of lycopene^15^, a biologically important carotenoid, derived from acetyl-CoA according to the Heimdallarchaeota MAGs (Fig. 1d, Supplementary Table 3). It has been mentioned that chlorophylls in photosynthesis ineffectively absorbed much light in the 450-550 nm (blue-green light) region of the solar radiation spectrum, while the light within this range can be effectively absorbed by carotenoids^9^. Moreover, carotenoids protect the organisms from photodamage by quenching both singlet or triplet states of bacteriochlorophylls under strong illumination and function as photosynthetic membrane stabilizers in chloroplasts^9^. Therefore, the biosynthesis of carotenoid in Heimdallarchaeota could coordinate with bacteriochlorophyll for high-efficiency photosynthesis, which provides an opportunity for competitive advantage in any particular habitat.

Besides photosynthetic pigments, many key factors related to reaction centers (including photosystem I (PS I) assembly proteins BtpA and Ycf3^16,17^; PSI subunit VII PsaC^18^; photosystem II (PS II) stability/assembly factor related protein Ycf48^19^; antenna proteins in PSII (phycocyanobilin lyase, CpcE and bilin biosynthesis protein, CpeU)^20,21^), carbon fixation system and ATP synthase have also been identified in Heimdallarchaeota MAGs (Extended Data Fig. 5, Supplementary Table 4). Taken together, the ubiquitous identification of photosynthetic apparatus and existence of both PS I and PS II in the genomes strongly suggest that Heimdallarchaeota are potential oxygenic photosynthetic organism^9^.

### Electron transfer and energy production in Heimdallarchaeota

Several lines of evidence support that Heimdallarchaeota perform photosynthesis based on our study. Logically, in the primary steps of photosynthesis, solar photons are absorbed by special membrane-associated pigment-protein complexes (light-harvesting antennas) and the electronic excitations are efficiently transferred to a reaction center^3,22^. The oxygen-evolving photosynthetic organisms have two photochemical reaction center complexes, PS I and PS II, that work together in a noncyclic electron transfer chain^22^. As a crucial electron carrier in PS I, phylloquinone was believed to be synthesized in Heimdallarchaeota (Fig. 2a, Supplementary Table 4). It is known that phylloquinone and its related compound menaquinone shared the same synthesis route for producing demethylphylloquinol^23^, which is further catalyzed to phylloquinone or menaquinone by a key enzyme named demethylphylloquinol methyltransferase (MenG) or demethylmenaquinone methyltransferase (UbiE)^23^. MenG is responsible for phylloquinone (vitamin K1) synthesis in both plants and Cyanobacteria, and UbiE is only in charge of menaquinone (vitamin K2) synthesis in microorganism^23^. Interestingly, both MenG and UbiE homologs were identified in Heimdallarchaeota MAGs (Supplementary Table 4). Consistently, in the phylogenetic tree, MenGs from photosynthetic organisms were clustered in a single branch which made a long distance from UbiE branches. While some MenG and UbiE homologs from plants, archaea together with Heimdallarchaeota were located exactly between these two branches (Fig. 2b, Supplementary Table 4). In addition, all enzymes catalyzing reactions to synthesize plastoquinol, which is further oxidized to plastoquinone by oxygen to phylloquinone (a key electron transporter in PS II), could also be identified in Heimdallarchaeota MAGs (Fig. 2c, Supplementary Table 4). Collectively, Heimdallarchaeota are believed to possess both PS I and PS II. Furthermore, subsequent electron acceptors including ferredoxin (Fd) and ferredoxin NADP^+^ reductase (FNR)^3^ were also found in Heimdallarchaeota MAGs (Fig. 3, Supplementary Table 4), which confers Heimdallarchaeota oxygenic photosynthetic capability through two photosystems as that in Cyanobacteria.

**Fig. 2.**
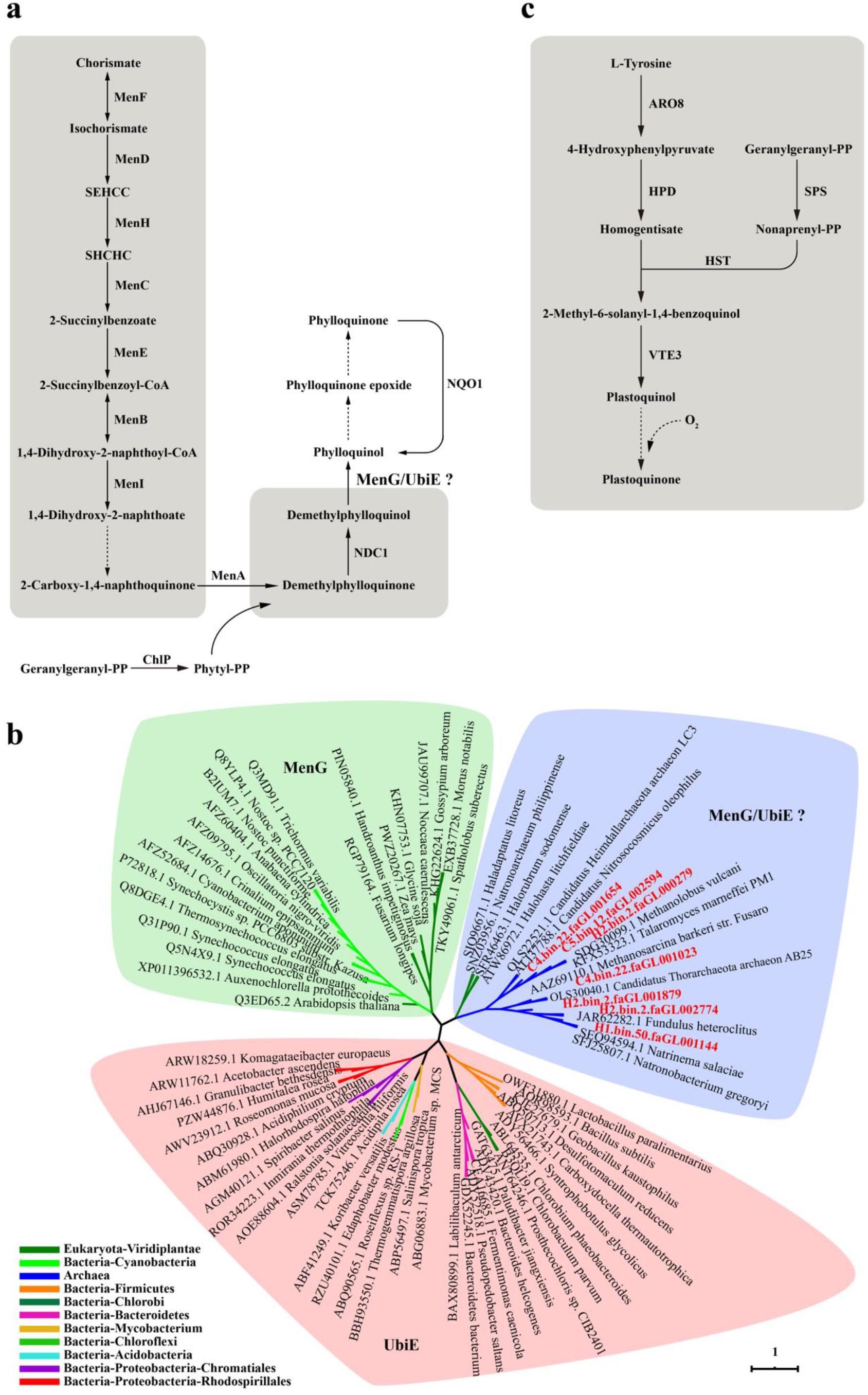
Phylogenomic analysis of phylloquinone and plastoquinone biosynthesis in Heimdallarchaeota. **a**, Analysis of phylloquinone biosynthesis in Heimdallarchaeota. The detail information of key enzymes involved in phylloquinone biosynthesis is listed in Supplementary Table 4. SEHCC, 2-Succinyl-5-enolpyruvyl-6-hydroxy-3-cyclohexene-1-carboxylate; SHCHC, (1R, 6R)-2-Succinyl-6-hydroxy-2,4-cyclohexadiene-1-carboxylate. **b**, Phylogenetic analysis of MenG/UbiE. An unrooted maximum-likelihood tree of MenG/UbiE homologs derived from different photosynthetic organisms identified in this work. The bootstrap support values 1,000. The green box highlights proteins annotated as MenG. The pink box highlights proteins annotated as UbiE. The blue box highlights proteins ambiguously annotated as MenG or UbiE. **c**, Analysis of plastoquinone biosynthesis in Heimdallarchaeota. ARO8, 2-aminoadipate transaminase. HPD, 4-hydroxyphenylpyruvate dioxygenase. SPS, All-trans-nonaprenyl-diphosphate synthase. HST, Homogentisate solanesyltransferase. VTE3, MPBQ/MSBQ methyltransferase. For panels a and c, the solid arrows indicate the enzymes associated with phylloquinone or plastoquinone biosynthesis identified in Heimdallarchaeota MAGs. Dotted arrows indicate the enzymes associated with phylloquinone or plastoquinone biosynthesis not identified in Heimdallarchaeota MAGs. The detail information of key enzymes involved in phylloquinone or plastoquinone biosynthesis is listed in Supplementary Tables 4 and 5.

**Fig. 3.**
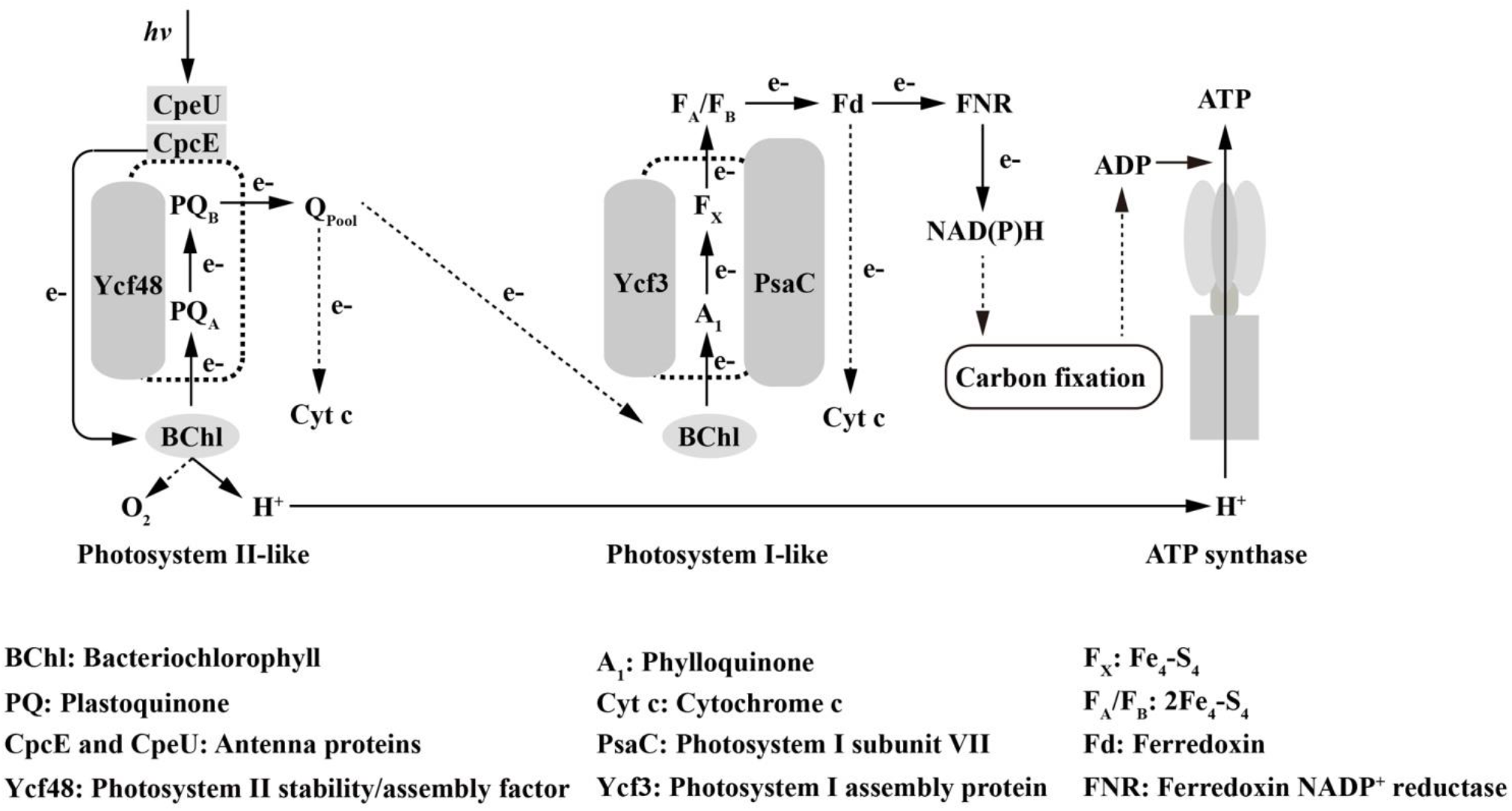
The inferred photosynthetic model of Heimdallarchaeota. Solid lines and arrows indicate the elements associated with photosystem and electron transport identified in Heimdallarchaeota MAGs, respectively. Dotted lines and arrows indicate the elements associated with photosystem and electron transport not identified in Heimdallarchaeota MAGs, respectively. The detail information of key enzymes involved in photosystem and electron transport is listed in Supplementary Table 4.

Overall, we proposed a complete oxygenic photosynthetic pathway existing in Heimdallarchaeota (Fig. 3). Firstly, PS II absorbs a photon of light to generate a redox-potential by antenna proteins and reduces plastoquinone as the terminal electron acceptor^3,22^. The electrons extracted from water are further transported via a quinone and the cytochrome c complex to PS I with the presence of complex III^24^ or alternative complex III (ACIII)^25^ or other homologs existing in Heimdallarchaeota. Meanwhile, electrons are removed from water by PS II, oxidizing it to molecular oxygen, which is released as a waste product. Then electrons are carried by phylloquinone and transferred to Fe_4_-S_4_ cluster to generate NADH with electrons from ferredoxin in PSI^3^. After the electron transfer, NAD(P)H is reduced with the electron delivering from ferredoxin^3,22^, which further participates in Calvin-Benson-Bassham cycle for carbon fixation and finally to synthesize ATP^3,22^.

### Heimdallarchaeota are photomixotrophic

Next, to gain further insight into the lifestyle of Heimdallarchaeota, we reconstructed a complete metabolic pattern according to our and published Heimdallarchaeota MAGs. Similar with other Asgard superphylum^26^, Heimdallarchaeota possess a mixotrophic lifestyle, which can simultaneously utilize the reverse tricarboxylic acid cycle (rTCA) for autotrophic CO_2_ assimilation^5,6^ and transport the exogenous organic matter through the metabolic circuitry for their catabolism (Fig. 4, Supplementary Table 5)^6^. For autotrophic metabolism, the Calvin-Benson-Bassham cycle was found to participate in carbon fixation through the RuBisCo and act as an intermediary between photosystems and ATP synthase for energy producing in Heimdallarchaeota (Fig. 4). Furthermore, a variety of polysaccharide-degrading enzymes, including chitinase, xylan/chitin deacetylase, diacetylchitobiose deacetylase and cellulase, were also found in Heimdallarchaeota by CAZy analysis (Extended Date Fig. 6 and Supplementary Table 5). These enzymes degraded polysaccharides like chitin, xylan and cellulose to produce oligosaccharides or monosaccharides for energy metabolism in Heimdallarchaeota. Moreover, methane metabolism pathway and Wood-Ljungdahl pathway were believed to exist in Heimdallarchaeota MAGs (Fig. 4).

**Fig. 4.**
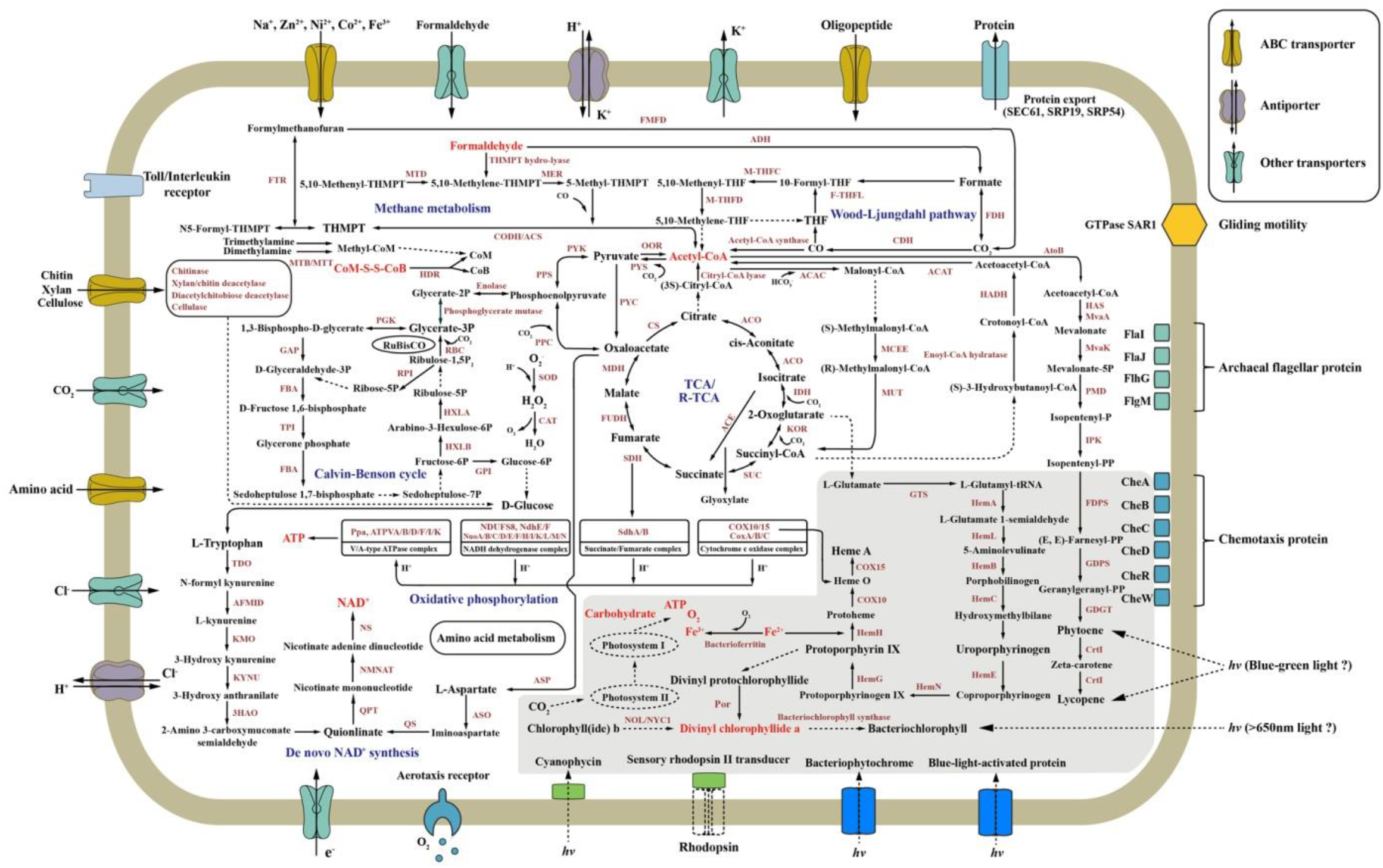
Reconstruction of photomixotrophic lifestyle in Heimdallarchaeota. Solid arrows indicate the enzymes associated with corresponding metabolic pathway identified in Heimdallarchaeota MAGs. Dotted arrows indicate the enzymes associated with corresponding metabolic pathway not identified in Heimdallarchaeota MAGs. The grey box highlights the photosynthetic pathway present in Heimdallarchaeota. The detail information of key enzymes mentioned in this figure is listed in Supplementary Table 5.

Notably, aerobic metabolic pathways were also found to be ubiquitous in Heimdallarchaeota^6,27^, which completely differed from metabolic characteristics of anaerobic Lokiarchaeota and Thorarchaeota^5,6,26,27^. Consistent with the results of previous studies^6^, the complete tricarboxylic-acid cycle (TCA) and oxidative phosphorylation pathway were found to support the aerobic respiration in Heimdallarchaeota. And both the aerobic kynurenine pathway and aspartate pathway for NAD^+^ *de novo* synthesis were also reconstructed in the present and previous published Heimdallarchaeota^6,28^. Particularly, the aerobic kynurenine pathway of NAD^+^ biosynthesis was exclusively found in Heimdallarchaeota compared with other archaea, which is only present in eukaryotes and very few bacterial groups^28^. This pathway has been considered to originate from the protoeukaryote ancestor in oxygen-containing niche, which might be acquired through horizontal gene transfer in Heimdallarchaeota^6^. In addition, other molecules in oxygen-dependent metabolism, such as aerotaxis receptor and bacterioferritin, were also identified in Heimdallarchaeota MAGs. All the above evidence indicates that oxygen must be present in the environment where Heimdallarchaeota inhabit. But this seems to contradict the strict anaerobic lifestyle existing in other Asgard archaea like Thorarchaeota and Lokiarchaeota in the same niche^5,6^. Therefore, we infer a more possible hypothesis that the oxygen presenting intracellular environment of Heimdallarchaeota might be generated by their performing photosynthesis. Consistently, Heimdallarchaeota have already evolved protection mechanisms against the formation of reactive oxygen species (ROS) via superoxide dismutase, catalase, and carotenoids (Fig. 4). Together, the metabolic reconstructions indicate that Heimdallarchaeota are photomixotrophic and have a facultative aerobic metabolism. If this is the fact, what is the ecological function or benefit for having the ability of photosynthesis in Heimdallarhaeota, which reside predominantly in marine sediments^5^? Recent study about rhodopsins identified in Heimdallarchaeota provides evidence of their light-exposed habitats, where Heimdallarchaeota could obtain enough energy from sunlight through photosynthesis^6^. The recovery of Heimdallarchaeota from deeper environments may be due to the high deposition rates characteristic for the sampling locations^6^. Nevertheless, there are plenty evidence showing that blue-green light with a wavelength range of 450-550 nm might exist in cold seep environment (~1100 m deep), and both long wavelength light (>650 nm) and (short wavelength light (<650 nm) could be detected in hydrothermal vents^29,30^, which provides necessary condition for photosynthetic process. The Heimdallarchaeota capable of detecting geothermal light and phototaxis could preferentially occupy an optimum habitat, which confer them evolutionary advantages in the competition for nutrient resources. Accordingly, a kind of cyanobacterium collects light and passes excitation energy uphill to the photochemically active pigments through longer-wavelength chlorophyll f, which facilitates this bacterium to survive in dark condition^31^. Thus, we suppose that possessing a presumptive photomixotrophic lifestyle may give Heimdallarchaeota more flexibility to survive or adapt to the deep-sea harsh conditions.

### A potential ancestor of eukaryotic photosynthetic organisms

Notably, given the close match of the emission spectra of geothermal light and the absorption spectra of bacteriochlorophylls *a* and *b*, photosynthesis was proposed to arise from bacteriochlorophyll *a*- or *b*-containing organisms near oceanic hydrothermal vents where weak infrared radiation could be detected^32^. In the evolution history of photosynthesis, the organisms were thought to initially use the bacteriochlorophyll pigments to sense infrared light, and started making use of the near-infrared part of sunlight when moving to shallow water through further adaptation of a primitive photosystem, and chlorophylls would be eventually developed to make use of higher energy (visible) light to split water^32,33^. Meanwhile, during the evolution process of oxygenic photosynthesis, a bioinorganic water-oxidizing complex (WOC) was thought to serves as a redox capacitor to accomplish the oxidation of two water molecules to produce O_2_ in PS II, which is comprised with a Mn_3_CaO_4_ distorted cubane structure bound to a fourth Mn by oxo-bridges^3,34^. And photoassembly of the WOC requires only Mn^2+^ and light to form the high-valent WOC^3^. Interestingly, an extremely high concentration of manganese element was detected in our sampling hydrothermal vent (Extended Data Fig. 2), which might promote WOC formation and play a key role in the acquiring and developing of photosynthesis in Heimdallarchaeota. Consistently, Heimdallarchaeota MAGs H2.bin.2 and H2.bin.81 derived from hydrothermal vent possess much more photosynthetic characteristics than those from cold seep in our present work. Therefore, in combination with the facts that Heimdallarchaeota are the most probable candidates for the archaeal protoeukaryote ancestor in previous report^6^ and the first identified photosynthetic archaea in this study, we propose that Heimdallarchaeota might be ancestor of eukaryotic photosynthetic organism.

The presence of oxygen-dependent pathways in Heimdallarchaea raises the possibility that the archaeal eukaryotic phtosynthetic ancestor could have also been a facultative aerobe, and the archaeal-photobacterial endosymbiosist gave birth to the eukaryotic phtosynthetic ancestor took place after the Great Oxidation Event^35^. Horizontal gene transfer is postulated to play a major role in the evolution of microbial phototrophs and that many of the essential components of photosynthesis have conducted horizontal gene transfer^22^. And Heimdallarchaea might obtain the photosynthetic apparatus through lateral gene transfer from their cyanobacterial endosymbionts^6,28^ because most of key factors associated with photosynthesis in Heimdallarchaea have high similarity with those from cyanobacteria (Supplementary Table 5). Our prediction is in agreement with the recent hypothesis that both the archaeal and bacterial eukaryotic ancestors have an oxygen-dependent metabolism^6^, in which the primordial function of the bacterial counterpart performing oxidative phosphorylation would not be detrimental to the existence of the archaeal host who exposed in oxygen environment. Even though the present results provide an updated perspective on the photosynthetic lifestyle of Heimdallarchaeota, further studies will be needed to elucidate the light utilizing strategies and evolutionary histories with their enrichment or even pure culture.

## Methods

### Sample collection and processing

Samples were collected from the cold seep in South China Sea and hydrothermal vent field in Okinawa trough (Expended Data Table 1) during the cruise of the R/V *Kexue* on July of 2017-2018. The sediment samples (C1, C4, C2 and C5) were collected from cold seep area in South China Sea at depth intervals of 0-20, 20-40, 40-60 and 280 cm. The other subsurface sediment (0-20 cm) samples (H1 and H2) were taken at the outside of the “black chimney” of the hydrothermal vent. Among the specimens, samples C1, H1 and H2 were collected through the Discovery remotely operated vehicle (ROV), sample C4 was obtained by the TV grab, while samples C2 and C5 were taken from the gravity sampler. Sediments were sealed into sterile sampling bags immediately after collection, and stored in −80 °C. DNA for metagenomics analysis was isolated from 20 g (wet weight) sediment per sample with the PowerSoil DNA Isolation Kit (Qiagen) following the manufacturer’s instructions.

### Analyses of environmental and chemical parameters of sampling sites

The temperature, salinity, underwater depth of sampling sites were recorded in real-time by SBE 25plus Sealogger CTD (SBE, USA), and concentrations of CO_2_ and CH_4_ of surface sediments were in situ measured with the CONTROS®HydroCO_2_ (CONTROS, Norway) and Hydro®CH_4_ (CONTROS, Norway), respectively. All these sensors were mounted on the Discovery ROV. For chemical element analyses, sediment samples from cold seep (C1, C2, C4 and C5) in South China Sea and hydrothermal vent field (H1 and H2) in Okinawa trough were dehydrated in an oven at 80 °C, respectively, until completely dry. After grinded, powder of samples were filtered through the 200-mesh screen. The obtained filtrate was further used to analyze the contents of different chemical elements, including Na, Mg, Fe, Cl, S, P, Mn, Zn, Ni and Co, by an S8 Tiger X-ray fluorescence spectrometry (BRUKER, Germany).

### Library construction and sequencing

DNA extracts were treated with DNase-free RNase to eliminate RNA contamination. Then the DNA concentration was measured by Qubit 3.0 fluorimeter (Thermo Fisher Scientific, USA). DNA integrity was evaluated by gel electrophoresis and 0.5 μg of each sample was used to prepare libraries. The DNA was sheared into fragments between 50-800 bp using Covaris E220 ultrasonicator (Covaris, UK). DNA fragments between 150 bp and 250 bp were secreted using AMPure XP beads (Agencourt, USA) and then were repaired using T4 DNA polymerase (ENZYMATICS, USA). These DNA fragments were ligated at both ends to T-tailed adapters and amplified for eight cycles. Finally, the amplification products were subjected to single-strand circular DNA libraries. All NGS libraries were sequenced on BGISEQ-500 platform (BGI, China) to obtain 100 bp paired-end raw reads. Quality control was performed by SOAPnuke (v1.5.6) (setting: −l 20 -q 0.2 -n 0.05 -Q 2 -d -c 0 −5 0 −7 1)^36^.

### Genomes assembly, binning and annotation

The raw shotgun sequencing metagenomic reads were dereplicated and trimmed by the BGI-Qingdao (BGI, China). The clean data were assembled using MEGAHIT (v1.1.3, setting: --min-count 2--k-min 33 --k-max 83 --k-step 10)^37^. Thereafter, metaBAT2^38^, Maxbin2^39^ and Concoct^40^ were used to automatically bin from assemblies. Finally, MetaWRAP^41^ was used to purify and generate data to get the final bins. Manual curation was adapted for reducing the genome contamination based on differential coverage, GC content, and the presence of duplicate genes. The completeness and contamination of the genomes within bins were then estimated by using CheckM^42^. Gene prediction for individual genomes was performed using Glimmer (v 3.02)^43^. The KEGG (Kyoto Encyclopedia of Genes and Genomes, Release 87.0), NR (Non-Redundant Protein Database databases, 20180814), Swiss-Prot (release-2017_07) and EggNOG (2015-10_4.5v) databases were used to annotate protein functions by default, and the best hits were chosen. Additionally, database of CAZy (Carbohydrate-Active enZYmes Database)^44^ was downloaded to search for carbohydrate active enzymes from genomic bins.

### Phylogenetic analyses

To reveal the phylum composition of assembled genomes in the archaea kingdom, the genomic sequences of Archaea were downloaded from NCBI ref genomes using Aspera (v3.9.8). Then, extract 37 marker genes in genomes (Supplemental Table 1) were chosen by Phylosift (v1.0.1)^45^ with automated setting. The concatenated sequences were trimmed using TrimAl (version 1.2)^46^ using gappyout function. Finally, maximum likelihood tree was calculated by using IQ-TREE (v1.6.12)^47^ with GTR+F+I+G4 model (-bb 1000) and shown by iTOL (v5)^48^. For phylogenetic analyses of protochlorophyllide reductase (Por), bacteriochlorophyll synthase (BCS) and demethylphylloquinol/demethylmenaquinone methyltransferase (MenG/ubiE) in Heimdallarchaeota bins, the related sequences were selected from archaea, bacteria and eukaryotes in NCBI and Swiss-Prot database. The Maximum-Likelihood phylogeny trees were constructed with WAG+G4, LG+F+I+G4 and LG+G4 model (-bb 1000) by using IQ-TREE, respectively, and showing with iTOL.

### Data availability

The Heimdallarchaeota genomic bins (C2.bin.3, C4.bin.14, C4.bin.22, C5.bin.12 and H2.bin.2) supporting the results of this study are available in NCBI Genbank under the accession numbers: SAMN13483368, SAMN13483369, SAMN13483392, SAMN13483370 and SAMN13483372 in BioProject PRJNA593668, respectively.

## Acknowledgements

This work was funded by the Strategic Priority Research Program of the Chinese Academy of Sciences (Grant No. XDA22050301), China Ocean Mineral Resources R&D Association Grant (Grant No. DY135-B2-14), National Key R and D Program of China (Grant No. 2018YFC0310800), the Taishan Young Scholar Program of Shandong Province (tsqn20161051), and Qingdao Innovation Leadership Program (Grant No. 18-1-2-7-zhc) for Chaomin Sun.

## Author Contributions

CS and RL conceived and designed the study. RL and JZ collected samples and supported the information of genomes. RL and RC analyzed the data. RL and CS wrote the manuscript with the input from all authors. All authors read and approved the final manuscript.

## Expanded data Figures

**Extended Data Fig. 1.**
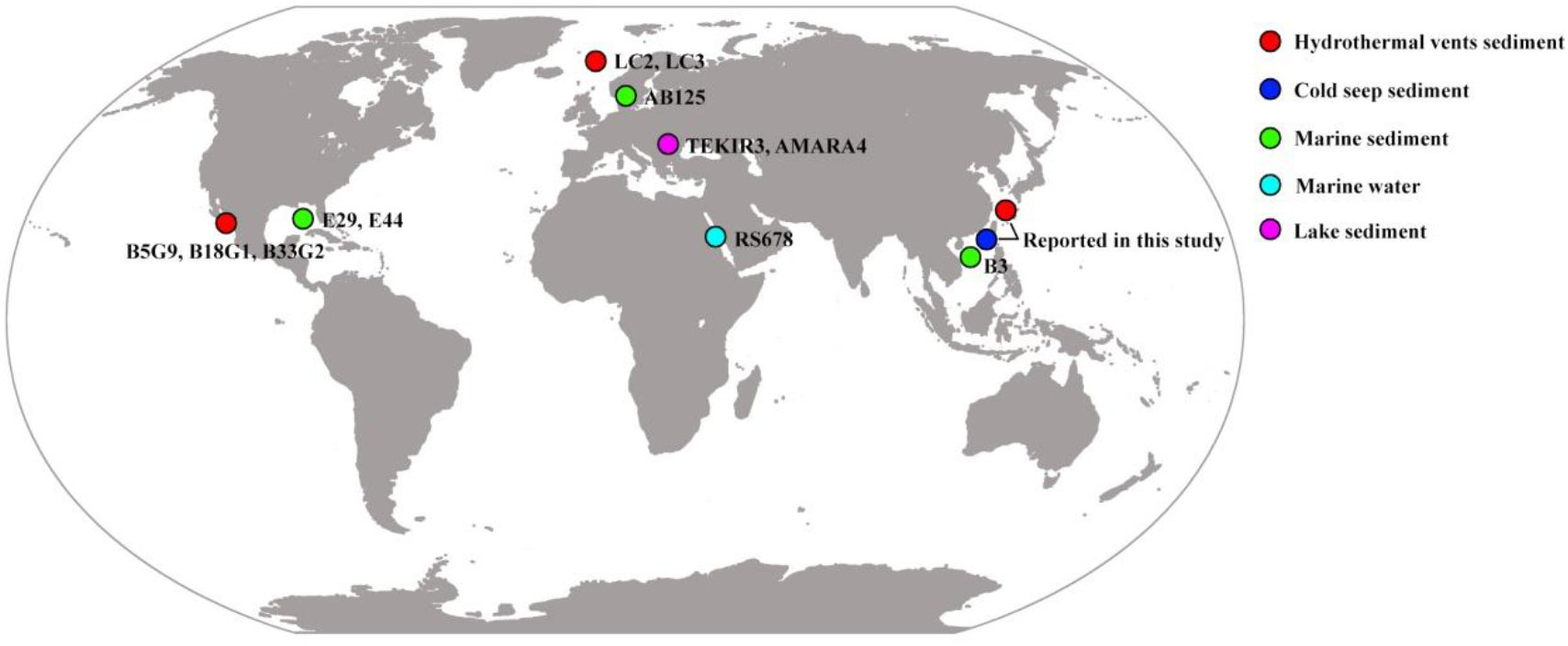
Global distribution of Heimdallarchaeota MAGs reported in previous and present studies. MAGs LC2 and LC3 are derived from Loki’s Castle. MAG AB125 is derived from Aarhus Bay (Denmark). MAG TEKIR3 is derived from Tekirghiol (Romania). MAG AMARA4 is derived from Amara (Romania). MAG RS678 is derived from Red Sea (Saudi Arabia). MAGs B5G9, B18G1 and B33G2 are derived from Guaymas Basin, Gulf of California (Mexico). MAGs E29 and E44 are derived from Atlantic Ocean. MAG B3 is derived from the north of the South China Sea (China).

**Extended Data Fig. 2.**
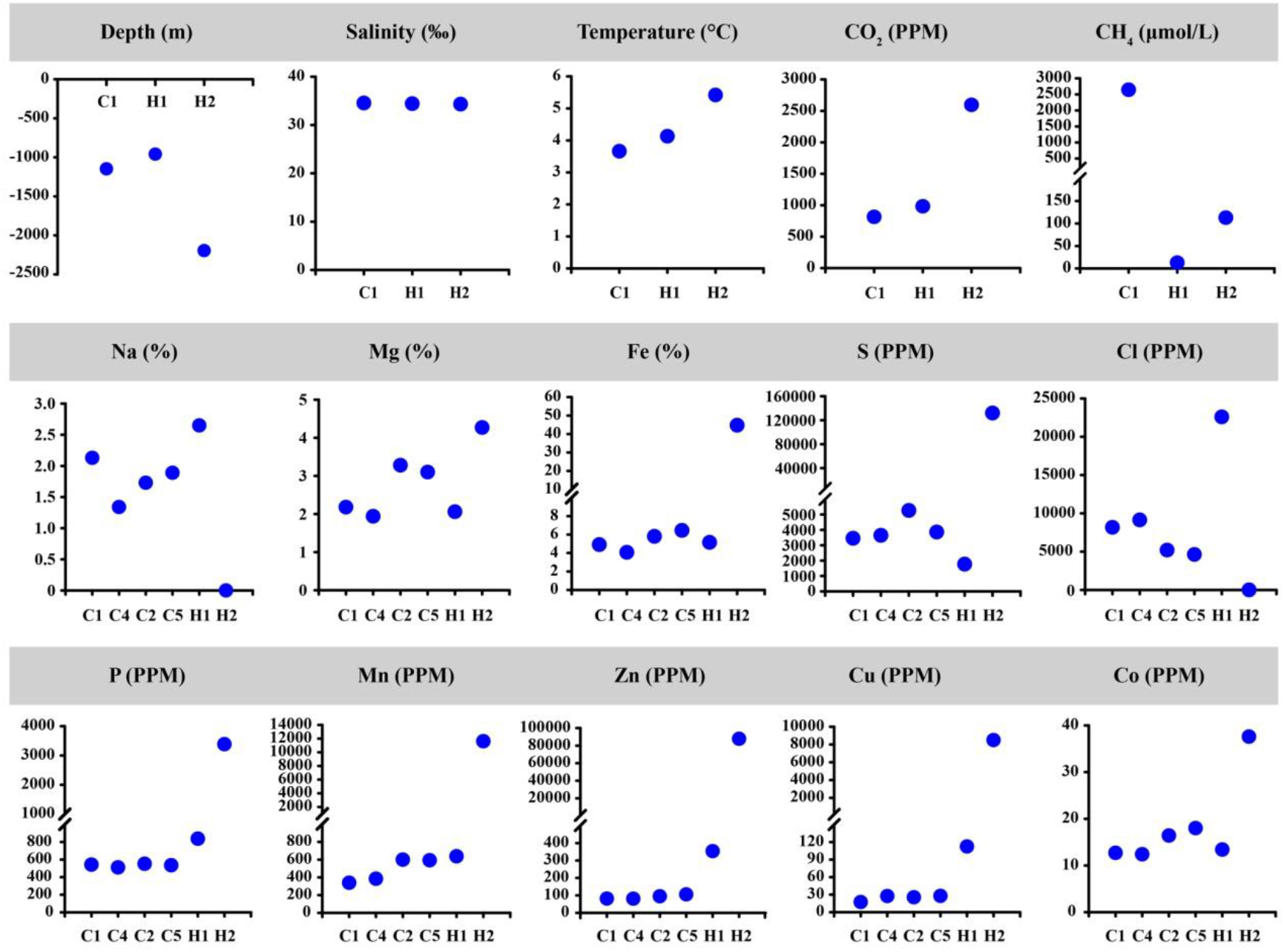
Analyses of environmental and chemical parameters of sampling sites in the deep-sea cold seep and hydrothermal vents. The temperature, salinity, underwater depth were recorded in real-time by SBE 25plus Sealogger CTD, and concentrations of CO_2_ and CH_4_ of surface sediments were in situ measured with the CONTROS®HydroCO_2_ and Hydro®CH_4_. Contents of different elements including Na, Mg, Fe, Cl, S, P, Mn, Zn, Ni and Co were measured by an S8 Tiger X-ray fluorescence spectrometry.

**Extended Data Fig. 3.**
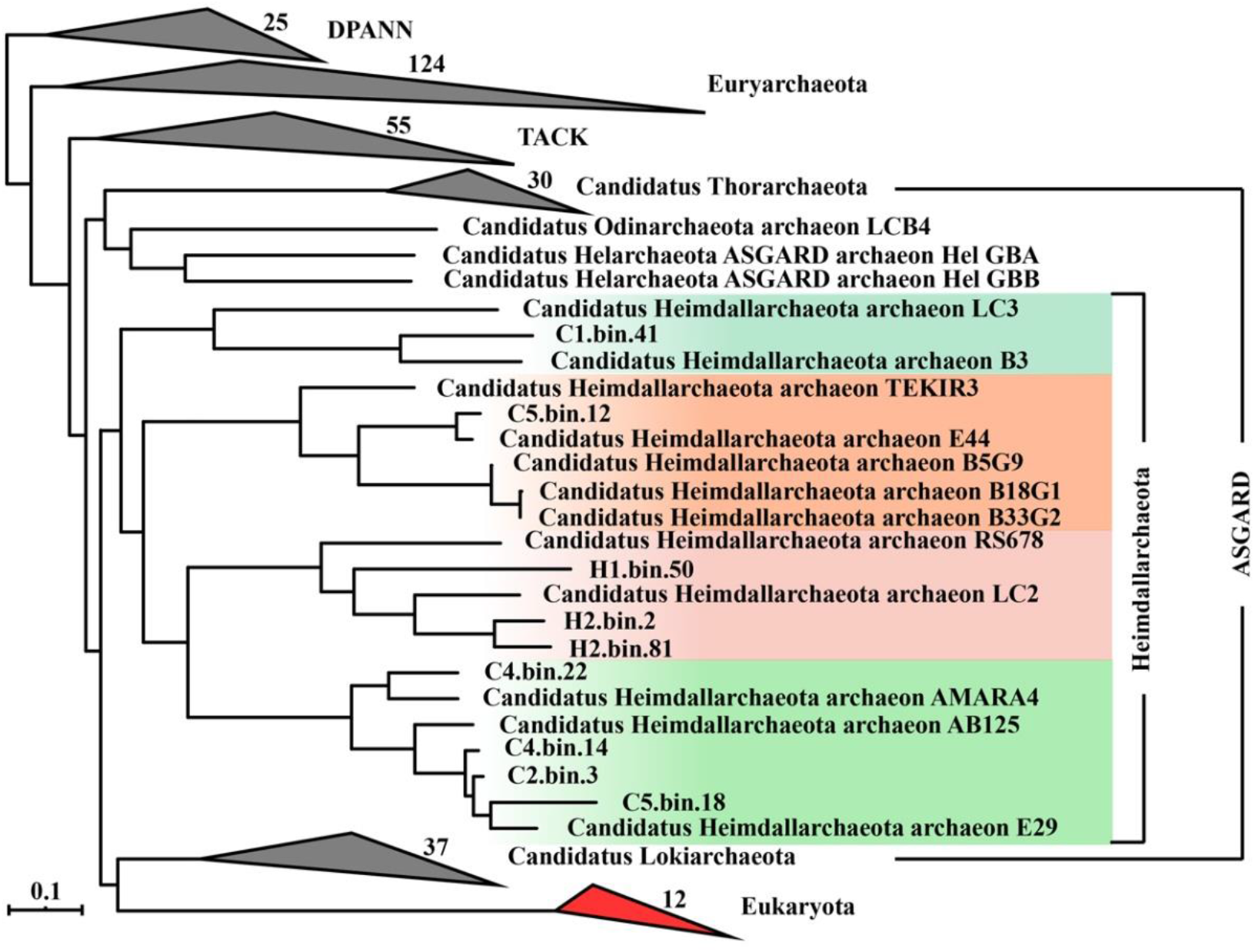
Maximum-likelihood phylogeny of superphyla Asgard, TACK, Euryarchaeota, DPANN and Eukaryota. Total 37 marker genes chosen by Phylosift, including 13 small subunit (SSU) and 16 large subunit (LSU) ribosomal RNA genes. The bootstrap support values 1000. All detailed sequence information of different species in compressed clades is listed in Supplementary Table 1.

**Extended Data Fig. 4.**
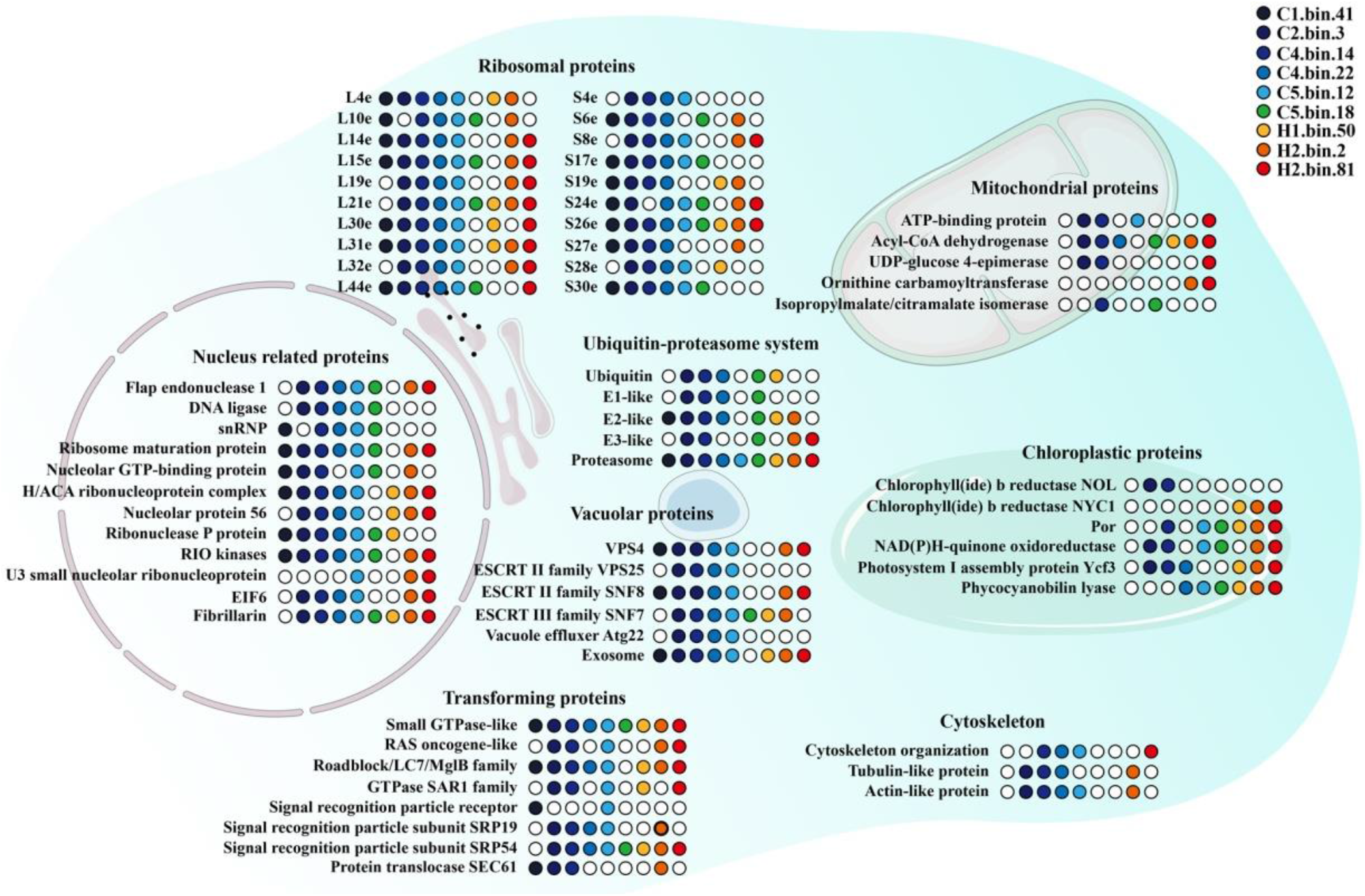
Eukaryotic signatures in Heimdallarchaeota. Schematic representation of a eukaryotic-like cell in which ESPs that have been identified in Heimdallarchaeota are highlighted. The overall illustration indicates that Heimdallarchaeota contain both reported eukaryotic signatures and unprecedented chloroplastic proteins. All detailed protein information mentioned in this figure is listed in Supplementary Table 2.

**Extended Data Fig. 5.**
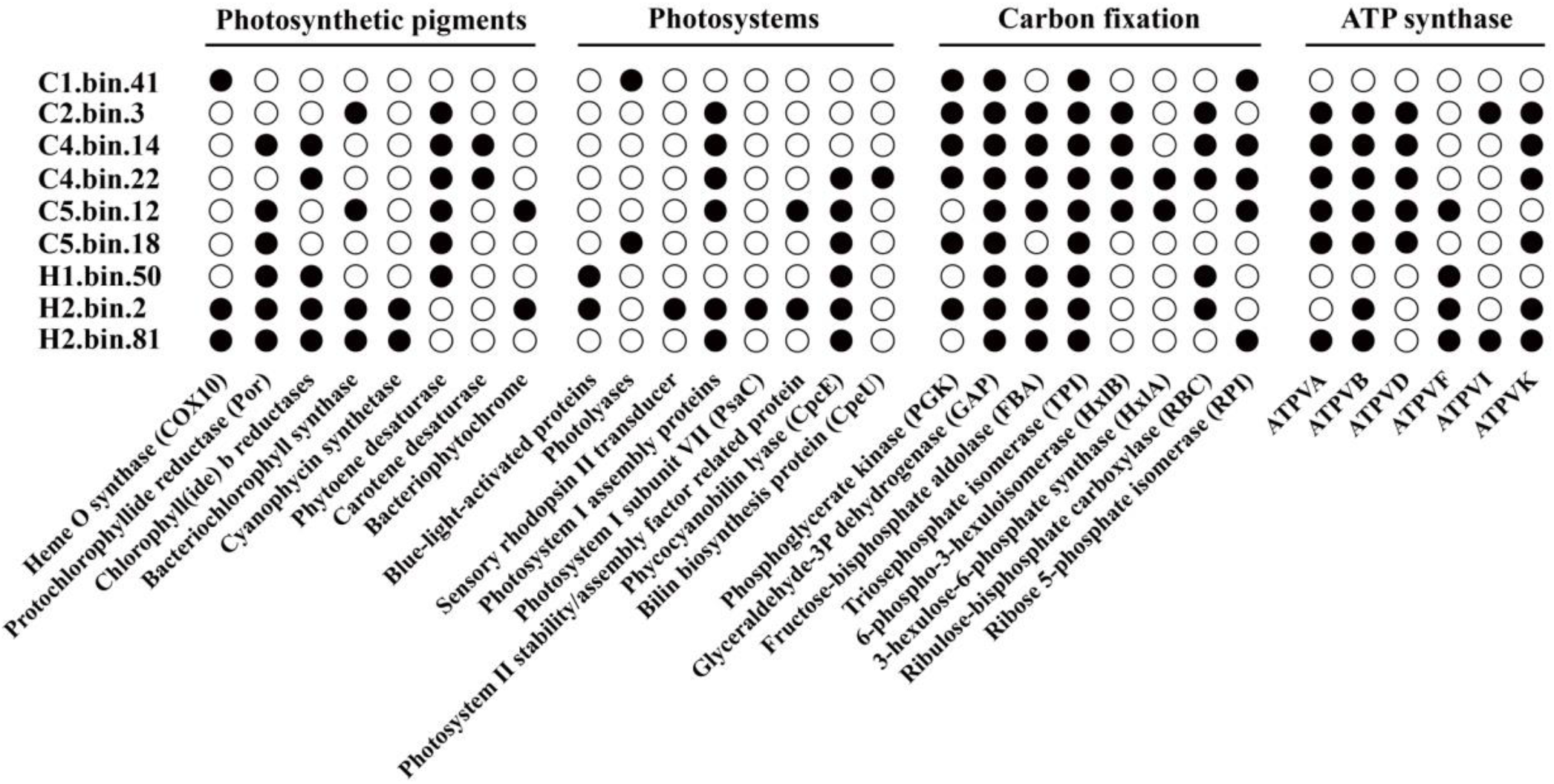
Photosynthetic apparatus identified in Heimdallarchaeota MAGs. All detailed protein information mentioned in this figure is listed in Supplementary Table 5. ATPVA~ATPVK, V/A-type H^+^/Na^+^-transporting ATPase subunits A~K.

**Extended Data Fig. 6.**
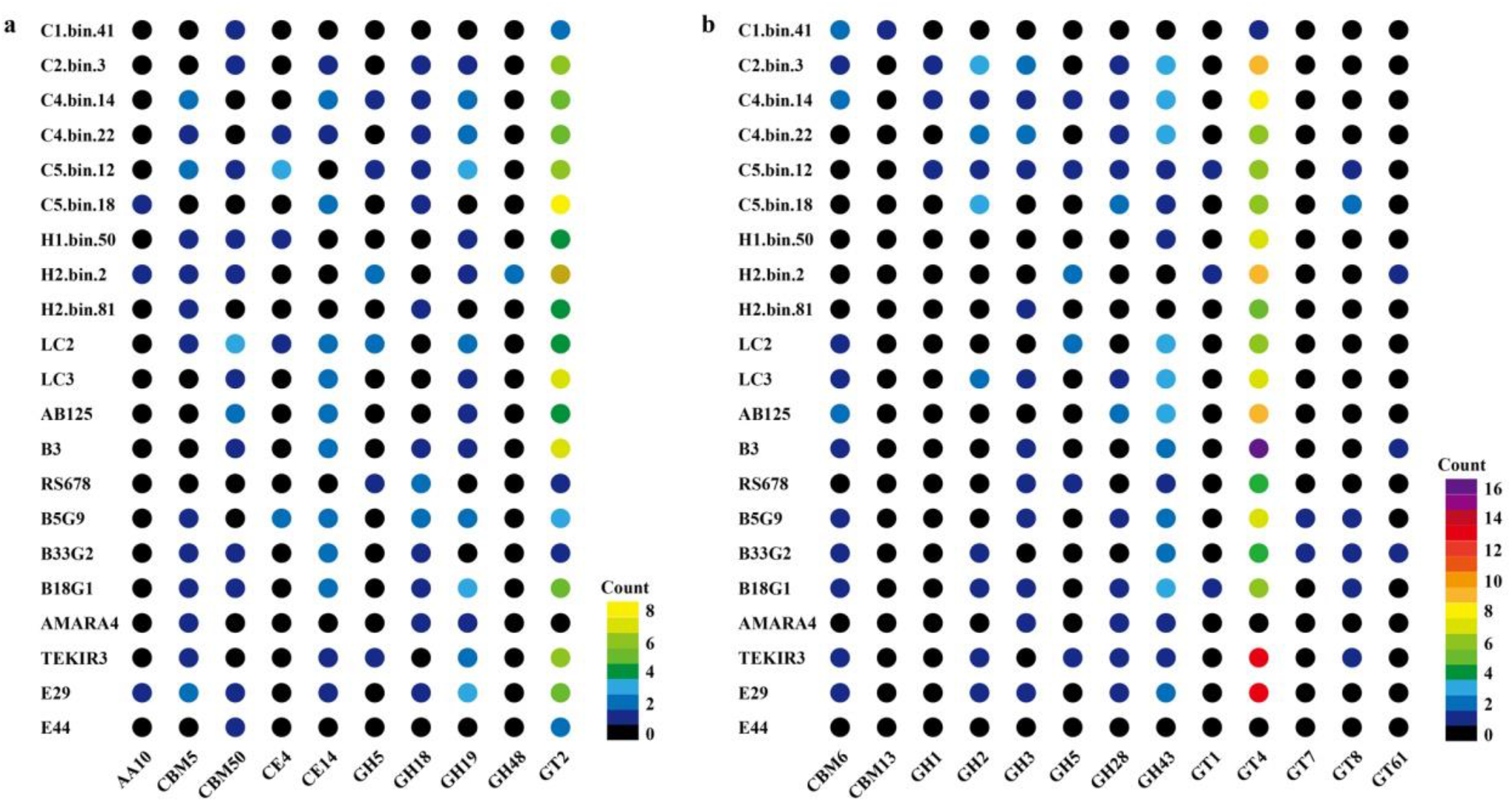
Chitin and xylan metabolic associated enzymes identified in Heimdallarchaeota MAGs by CAZy analysis. **a**, Chitin metabolic related enzymes identified in Heimdallarchaeota MAGs. **b**, Xylan metabolic associated enzymes identified in Heimdallarchaeota MAGs. AAs, auxiliary activities. CBMs, carbohydrate-binding modules. CEs, carbohydrate esterases. GHs, glycoside hydrolases. GTs, glycosyltransferases. All detailed protein information mentioned in this figure is listed in Supplementary Table 5.

