## Extended Data Tables for "Heimdallarchaeota harness light energy through photosynthesis"

**Expanded Table 1 Location information of samples**

| Sample name | Sample type | Location | Longitude and latitude | Depth (m) |
| --- | --- | --- | --- | --- |
| C1 | Sediment | Cold seep | E 119º17'09.106''  N 22º06'55.114'' | 1146 |
| C4 | Sediment | Cold seep | E 119º17'06.436''  N 22º06'55.169'' | 1137 |
| C2, C5 | Sediment | Cold seep | E 119º17'07.322''  N 22º06'58.598'' | 1121 |
| H1 | Sediment | Hydrothermal vent | E 126º53'51.85''  N 27º47'11.26'' | 958 |
| H2 | Sediment | Hydrothermal vent | E 124º22'23.374''  N 25º15'49.868'' | 2194 |

**Expended Data Table 2 Assembly statistics and quality metrics of reconstructed Heimdallarchaeota genome bins**

| Bin name | C1.bin.41 | C2.bin.3 | C4.bin.14 | C4.bin.22 | C5.bin.12 | C5.bin.18 | H1.bin.50 | H2.bin.2 | H2.bin.81 |
| --- | --- | --- | --- | --- | --- | --- | --- | --- | --- |
| Completeness (%) | 54.84 | 86.84 | 91.58 | 92.52 | 80.84 | 52.90 | 52.02 | 78.03 | 51.03 |
| Contamination (%) | 4.672 | 5.607 | 3.738 | 3.738 | 6.540 | 7.512 | 3.342 | 5.295 | 1.869 |
| GC (%) | 35.30 | 33.50 | 33.40 | 34.20 | 33.80 | 33.60 | 36.50 | 34.30 | 34.40 |
| N50 (bp) | 8641 | 10642 | 41762 | 25712 | 11218 | 2402 | 1661 | 15481 | 14781 |
| Size (Mbp) | 1.06 | 3.06 | 3.27 | 2.82 | 3.24 | 2.12 | 1.99 | 4.85 | 2.80 |
| Number of genes | 1079 | 1178 | 3084 | 2593 | 3128 | 2187 | 2644 | 1215 | 2637 |
